# PROPERMAB: an integrative framework for *in silico* prediction of antibody developability using machine learning

**DOI:** 10.1101/2024.10.10.616558

**Authors:** Bian Li, Shukun Luo, Wenhua Wang, Jiahui Xu, Dingjiang Liu, Mohammed Shameem, John Mattila, Matthew Franklin, Peter G. Hawkins, Gurinder S. Atwal

**Affiliations:** Regeneron Pharmaceuticals, 777 Old Saw Mill River Road, Tarrytown, NY 10591, USA

**Keywords:** antibody developability, machine learning, in silico prediction, feature engineering

## Abstract

Selection of lead therapeutic molecules is often driven predominantly by pharmacological efficacy and safety. Candidate developability, such as biophysical properties that affect the formulation of the molecule into a product, is usually evaluated only toward the end of the drug development pipeline. The ability to evaluate developability properties early in the process of antibody therapeutic development could accelerate the timeline from discovery to clinic and save considerable resources. *In silico* predictive approaches, such as machine learning models, which map molecules to predictions of developability properties could offer a cost-effective and high-throughput alternative to experiments for antibody developability assessment. We developed a computational framework, P_ROPERMAB_, for large-scale and efficient *in silico* prediction of developability properties for monoclonal antibodies, using custom molecular features and machine learning modeling. We demonstrate the power of P_ROPERMAB_ by using it to develop models to predict antibody hydrophobic interaction chromatography retention time and high-concentration viscosity. We further show that structure-derived features can be rapidly and accurately predicted directly from sequences by pre-training simple models for molecular features, thus providing the ability to scale these approaches to repertoire-scale sequence datasets.

## Introduction

Monoclonal antibody (mAb)-based biologics have become a major therapeutic modality in recent years [1, 2]. As of December 2023, more than 100 antibody therapeutics have been marketed and the number of antibody therapeutics in late-stage clinical studies has surpassed 130 [3]. For several disease areas such as oncology [4, 5], inflammation, and infectious disease [6], antibodies have become the predominant treatment options [1].

Despite the continuous advances in antibody discovery and development technologies, bringing an antibody from discovery to a marketed product remains a costly process with a high attrition rate [7]. While attrition may be caused by various efficacy and safety reasons, in the early-development stage poor developability properties pose unique challenges in product formulation and Chemistry, Manufacturing, and Control (CMC) process development [7–9]. A successful antibody drug development program relies on the selection of a lead candidate that is developable, with preferred properties in expression titer, purification yield, solubility, viscosity, stability and administration compatibility. Poor developability properties may add considerable cost to development and slow the timeline from discovery to clinic. Some developability issues such as high viscosity may be mitigated by optimization of excipients [10, 11] albeit often resulting in a deviation from platform formulation and manufacturing process. Other developability issues such as increased hydrophobicity or poor stability may require extensive optimization of purification process or even engineering of the sequence [12]. Given the considerable time and cost associated with off-platform manufacturing process and formulation or failure of development, it is essential to prioritize sequences that are less likely to pose developability issues when candidates are selected [7, 8].

In early-stage screening, candidate molecules are typically selected from a library of well over a thousand antibodies. Experimental assessment of the developability properties of such a library is challenging if not infeasible. This is in part due to the requirement for sufficient purified materials for each antibody and/or the development of high-throughput assays [13, 14]. As an alternative, computational methodologies for developability assessment are gaining increased attention [15] and several computational strategies have recently been introduced for predicting antibody solubility [16, 17], aggregation [18–20], thermostability [21], hydrophobic interaction chromatography (HIC) retention time [20, 22], pharmacokinetics [20, 23, 24], and high-concentration viscosity [20, 23, 25–28].

Historically, *in silico* developability assessment strategies have typically been heuristics that are based on individual molecular features designed to associate with a specific developability attribute [29, 30] or simple linear statistical or empirical models that consider only a handful of features [31, 32]. More recently, machine-learning (ML) models have been increasingly used for *in silico* developability assessment [14, 16, 21–23, 26, 27]. In addition to larger and higher-quality experimentally derived training data, having informative features as input is critical to the predictive performance of ML models. Several commercial software packages are available and can calculate a large set of input features. However, users are typically limited to a set of predefined features and implementing novel, user-designed features can be challenging in part due to the lack of source code access. Moreover, these software packages are typically not integrated with the Python ML ecosystem, thus including them into automated, Python-based ML model development workflows can be challenging.

Here, thanks to recent advances in 3D structure prediction and sequence modeling for proteins [33–35] and antibodies [36, 37], we developed a versatile and integrative computational tool called P_ROPERMAB_ to enable *in silico* developability assessment of mAbs using machine learning. This is accomplished by P_ROPERMAB_’s ability to efficiently calculate a diverse set of features, ease of engineering new features, and through its seamless integration with the Python ML ecosystem. We show the practical utility of P_ROPERMAB_ by applying it to predict mAb HIC retention time and high-concentration viscosity and demonstrate a computationally efficient and practically effective approach for applying P_ROPERMAB_ to repertoire-scale sequence datasets.

## Results

### Design and overview of P_ROPERMAB_

We developed P_ROPERMAB_ to facilitate *in silico* prediction of mAb developability properties with the only assumption that mAb sequences are available at the point of assessment. The design of P_ROPERMAB_ was guided by the preference for integrating features at both sequence and structure levels, ease of engineering new features, and the need for robust model validation, especially in the data-limited, supervised learning regime such as antibody developability prediction. At a high level, given the pair of sequences of the heavy and light chains of a mAb, one can use P_ROPERMAB_ to calculate a predefined set of sequence-based features (the Sequence Featurizer component in Figure 1, and Supplementary Table 1), extract sequence embeddings from pretrained protein language models [34, 37] (not shown in Figure 1), employ a deep learning-based protein 3D structure prediction method to predict the 3D structure for the Fv domain [36] (the Structure Predictor component in Figure 1), and calculate a predefined set of structure-based features from the predicted structures (the Structure Featurizer component in Figure 1, and Supplementary Table 2). Once features have been calculated both supervised and unsupervised learning can be pursued depending on the availability of experimentally measured developability data and the questions being asked. To facilitate the development of supervised models, we designed a ModelTrainer class that provides a unified interface for model training and validation (the Model Trainer component in Figure 1). The ModelTrainer class interfaces with the scikit-learn package [38] and encapsulates robust cross validation strategies for model parameter fitting, hyper-parameter tuning, and performance estimation.

**Figure 1.**
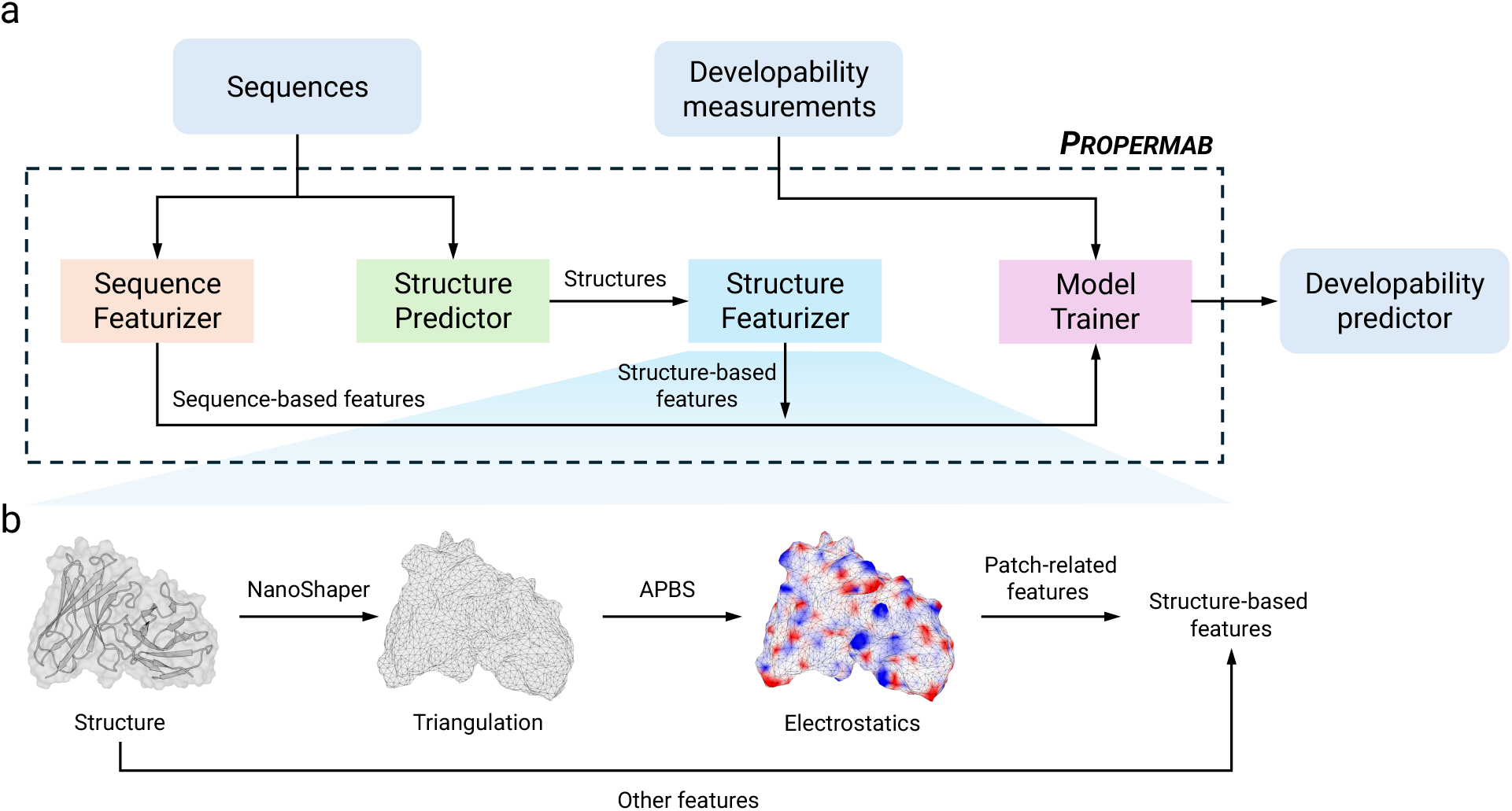
A schematic of P_ROPERMAB_ highlighting its main components. The main components of P_ROPERMAB_ were designed to facilitate feature calculation and supervised ML model development. Given antibody sequences, one uses the Sequence Featurizer component to calculate sequence-based features. To calculate structure-based features, one first uses the Structure Predictor component to predict 3D structures, which are then used as input to the Structure Featurizer component for feature calculation. Both the Sequence and Structure Featurizers currently offer a diverse set of features and can be easily extended with new, custom-designed features. Once features have been calculated, the training and validation of ML models are done through the Model Trainer component.

P_ROPERMAB_ is open source and implemented purely in Python. As such, it offers algorithmic transparency, ease of engineering new features, and is seamlessly integrated with other Python-based, open-source ecosystems for scientific computing such as machine learning and bioinformatics. In the following sections, we first demonstrate the ease of engineering new features in P_ROPERMAB_ by showcasing the implementation of a set of novel structure-based features that, to our knowledge, have not previously been used in mAb developability prediction. We then discuss examples of using P_ROPERMAB_ to develop ML models for predicting mAb HIC retention time and high-concentration viscosity, two important properties considered during mAb therapeutic product and process development. Finally, because predicting 3D structures and calculating structure-based features can be computationally costly for large, multi-repertoire-scale datasets, we demonstrate that one can train simple regularized linear models to accurately predict structure-based features directly from sequence. Using predicted features as input results in only a slight drop in performance compared with using features calculated from structures, suggesting a cost-effective approach for large datasets.

### P_ROPERMAB_ implements a diverse set of molecular features and offers the ease for engineering new features

P_ROPERMAB_ currently implements a diverse set of molecular features that are either derived from sequences (8 features, Supplementary Table 1) or structures (27 features, Supplementary Table 2). These features characterize the biophysical attributes of antibodies, such as electrostatic charge distribution, hydrophobicity, and solvent exposure, either based on amino acid sequences or the 3D structures of the Fv domain. For several charge and hydrophobicity related features, P_ROPERMAB_ also offers versions specific to or near the complementarity determining regions (CDRs) because they play the central role in antibody-antigen interaction. These features were largely inspired by current understanding of the molecular and biophysical mechanisms contributing to poor solution and colloidal properties and the fact that some of them have been used in the existing literature [14, 24, 39–41]. For example, the utility of surface patches (Figure 1, Methods, and Supplementary Table 2) in antibody developability assessment has been demonstrated in several recent studies [20, 24, 40, 41].

Out of the current set of features computed by P_ROPERMAB_, several of the sequence-derived, charge- or hydrophobicity-related features have been previously reported and used for *in silico* developability assessment [22, 27]. However, P_ROPERMAB_ offers the flexibility to easily customize some of the previously reported features for specific antibody regions. For example, we customized the net_charge feature for the CDRs (net_charge_cdr) and solvent-exposed residues (exposed_net_charge). Because P_ROPERMAB_ implements a triangle mesh-based strategy to compute surface patches, the vertices of the triangle mesh can be assigned with various biophysical properties and can also be mapped to residues (Figure 1 and Methods). Such information can be leveraged to create customized features such as number of patches and largest patch area of various physicochemical nature (i.e. positively charged, negatively charged, hydrophobic) and for specific antibody regions. More importantly, P_ROPERMAB_ also enables the design of novel features to leverage expert domain knowledge on the developability property of interest. For example, hypothesizing that the spatial pattern of biophysical features affects inter-molecular interactions, we implemented a variant of the Ripley’s K statistic [42, 43] for charged and aromatic residues (Figure 2 and Supplementary Table 2) as features for developability assessment. The Ripley’s K is a spatial statistic that characterizes spatial correlation of point patterns and is defined as the fraction of point pairs for which the distance is less than a distance cutoff (Figures 2a and 2b). Our implementations of the Ripley’s K statistics characterize the clustering or dispersion of charge and aromatic features over the surface of antibody Fv domain at a given distance threshold (Figure 2c). As discussed in the next two sections, some of the Ripley’s K features are proved to be informative in predicting antibody HIC retention time and viscosity.

**Figure 2.**
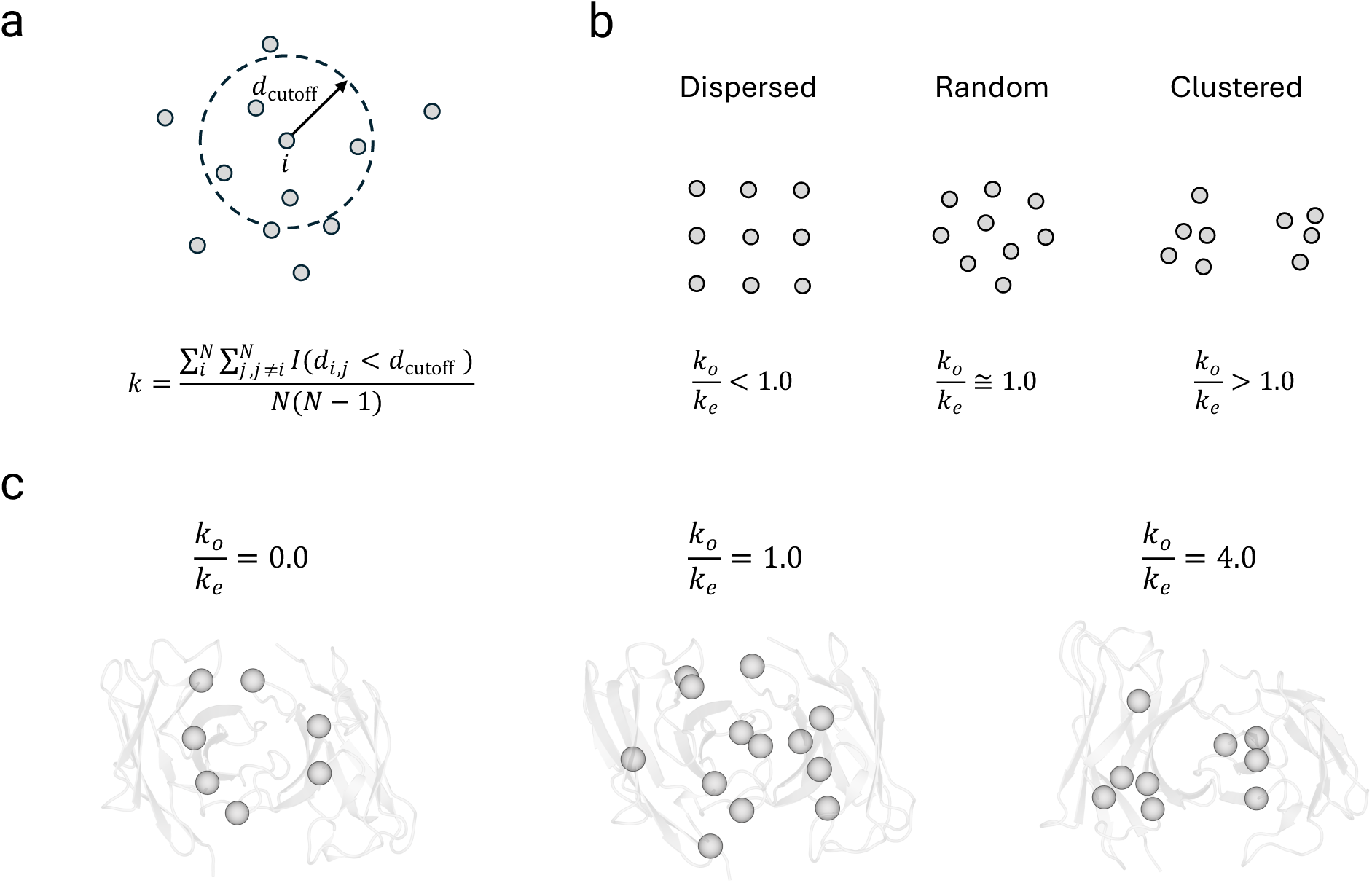
Implementation of a variant of the Ripley’s K function as molecular features. **a**) An illustration of the computation of the K function at contact distance cutoff *d*_cutoff_. In the formula, *N* is the total number of points, *d*_*i,j*_ is the Euclidean distance between point *i* and point *j*, *I* is the indicator function that evaluates to 1 when *d*_*i,j*_ is less than *d*_cutoff_ and 0 otherwise. Thus, *k* is the fraction of point pairs for which the distance is less than *d*_cutoff_. **b**) Schematics showing three different spatial patterns of point clouds. Here, *k*_*o*_ is the Ripley’s K based on the observed pattern, *k*_*e*_ is the expected Ripley’s K when the points are randomly scattered. Note that the *d*_cutoff_ in this panel much smaller than the *d*_cutoff_ in panel a). **c**) Examples of the Ripley’s K spatial statistic feature for surface aromatic residues. Here, only the Cα atoms of solvent-exposed aromatic residues are shown. Distances are measured between pairs of Cα and *d*_cutoff_ is set to 8 Å.

### Application of P_ROPERMAB_ to predict HIC retention time

We developed P_ROPERMAB_ as a general framework for feature engineering and ML model development for mAb developability prediction. As examples of use cases, we first demonstrate P_ROPERMAB_’s utility by applying it to predict the hydrophobic interaction chromatography retention time (HIC RT) and in the next section we apply it to predict high-concentration viscosity.

When screening for candidate molecules, it is important to consider a molecule’s hydrophobicity profile because increased hydrophobicity of mAbs often contributes to higher likelihood of precipitation and aggregation [41, 44], reduced purification yield or off-platform purification process. Hydrophobic interaction chromatography (HIC) is a standard biophysical method commonly used in protein purification [45] and HIC RT is characteristic of individual molecule’s surface hydrophobicity with longer RTs correlating with higher degree of hydrophobicity [46]. To demonstrate the utility of P_ROPERMAB_, we computed all 35 features for 135 mAbs for which the HIC RTs have been consistently measured [47] and first analyzed the correlations of individual features with HIC RT (Figure 3a). To account for the effect of 3D conformation on feature values (Supplementary Figure 1 and Supplementary Table 3), we computed structure-based features across five predicted structures and took the averages. Our analysis shows that about two thirds of the features (23 out of 35) are significantly correlated with HIC RT (Figures 3a) among which hyd_patch_area_cdr (total area of hydrophobic patches near CDRs, Methods) has the highest correlation, suggesting that mAbs likely bind the stationary phase of the HIC column through hydrophobic patches on their CDRs. In addition to hyd_patch_area_cdr, our analysis shows that the aromatic_asa feature (aromatic surface area) also has a strong correlation with HIC RT (Figures 3a and 3c). Interestingly, aromatic residues have previously been found to be enriched in aggregation prone regions in mAbs [48]. We also note that two novel, Ripley’s K derived features, neg_ripley_k (Ripley’s K ratio for negatively charged residues) and aromatic_ripley_k (Ripley’s K ratio for aromatic residues), are also found to be significantly correlated with HIC RT (Figure 3a and 3d). While the correlations are only moderate, it nevertheless demonstrates that one can engineer rather sophisticated and predictive features using our framework.

**Figure 3.**
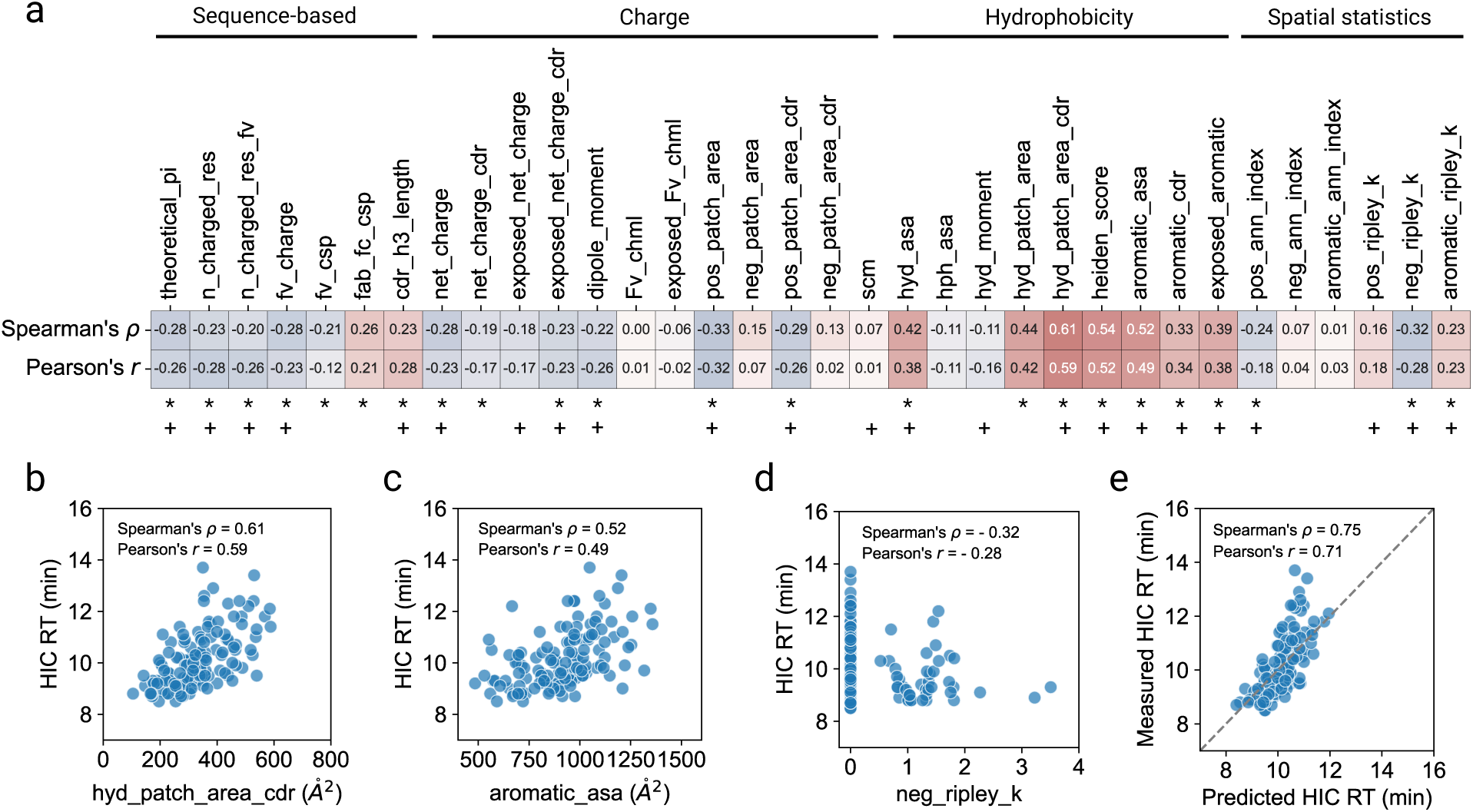
Applying P_ROPERMAB_ to predict mAb HIC retention time (HIC RT). **a**) Spearman’s ρ and Pearson’s *r* correlation coefficients of each feature with the HIC RTs of 135 mAbs from [47]. We excluded the two mAbs for which the HIC RTs were not determined but arbitrarily set to 25 min. The fc_charge feature is omitted because these molecules have identical constant region sequences [47], which resulted in identical values of the fc_charge feature. Features with a statistically significant (*p* < 0.05, adjusted via the Benjamini-Hochberg procedure) [49] Spearman’s ρ are indicated with a * symbol. **b, c, d**) Scatter plots of HIC RT vs. a few examples of predictive features. The hyd_patch_area_cdr feature (total hydrophobic patch area near CDRs) has the highest correlation with HIC RT, which is followed by the heiden_score and aromatic_asa feature (total aromatic solvent accessible surface area). The spatial statistic neg_ripley_k, which was designed to characterize the clustering or dispersion of negatively charged residues on antibody surface, has a moderate but significant negative correlation (Spearman’s ρ = −0.32, adjusted *p* < 0.05) with HIC RT. A neg_ripley_k of 0 indicates that there was no pair of negatively charged residues that are less than 8 Å from each other. **e**) Leave-one-out cross-validation results of an ElasticNet regressor trained to predict mAb HIC RT using all the features. Features with a non-zero coefficient in the ElasticNet model are indicated with a + symbol in panel **a**).

Encouraged by the correlations of features with HIC RT, we then trained an ElasticNet model using all molecular features currently available in P_ROPERMAB_ as input to predict HIC RT. We used a nested cross-validation strategy to estimate the performance of the model (Methods). Our nested leave-one-out cross-validation (LOOCV) evaluation shows that the ElasticNet regression model predicts HIC RT with a Pearson’s *r* = 0.71 and a Spearman’s ρ = 0.75, substantially better than any of the individual features (Figure 3e). We note that due to the small size of the dataset, we opted not to do explicit feature selection. However, after inspecting the model coefficient associated with each feature, we found that the set of features with a non-zero coefficient in the ElasticNet model is highly concordant with the set of features that are significantly correlated with HIC RT (Figure 3a).

### Application of P_ROPERMAB_ to predict high-concentration viscosity

Viscosity is a key developability attribute that can significantly affect both manufacturing and subcutaneous injection. mAb solutions with high viscosities add challenges to manufacturing, delivery, and administering [50]. To demonstrate the utility of P_ROPERMAB_ in predicting viscosity, we applied it to an internal viscosity dataset consisting of 58 unique IgG4 mAbs and two previously published datasets of the IgG1 isotype [25, 51] whose viscosities have been measured at 150 mg/mL. To our knowledge, our IgG4 dataset is the largest dataset of measured viscosities of the IgG4 isotype reported at the time of writing. As with HIC RT, our analysis shows that several features are strongly correlated with viscosity in the two previously published datasets (Supplementary Figure 2). However, no single feature is strongly correlated with viscosity in our internal IgG4 dataset (Figure 4a). This discrepancy is likely due to the fact that the two previously published datasets are of the IgG1 isotype and that molecules in one of the dataset, PDGF38, were derived from the same parent molecule and designed to optimize surface electrostatic properties for better viscosity profile [51]. Nevertheless, several features characterizing charge asymmetry across the VH and VL domains such as dipole_moment and Fv_chml are found to be at least weakly correlated with viscosity in our IgG4 dataset (Figures 4a, 4b and 4c). It has been previously shown in a much smaller dataset (14 IgG4 mAbs) that the spatial charge map (SCM) score [29] can correctly classify IgG4 mAbs into low or high viscosity with ∼80% accuracy [52]. Interestingly, this finding is not supported by our analysis of a much larger dataset because we find almost no correlation between the SCM score and viscosity (Spearman’s ρ = 0.12 and a Pearson’s *r* = 0.07) (Figures 4a and 4d). While the molecular origins of viscosity behaviors of mAb solutions are believed to be quite complex [53], our findings with a larger dataset provides strong support for the role that charge asymmetry plays in IgG4 mAb viscosity.

**Figure 4.**
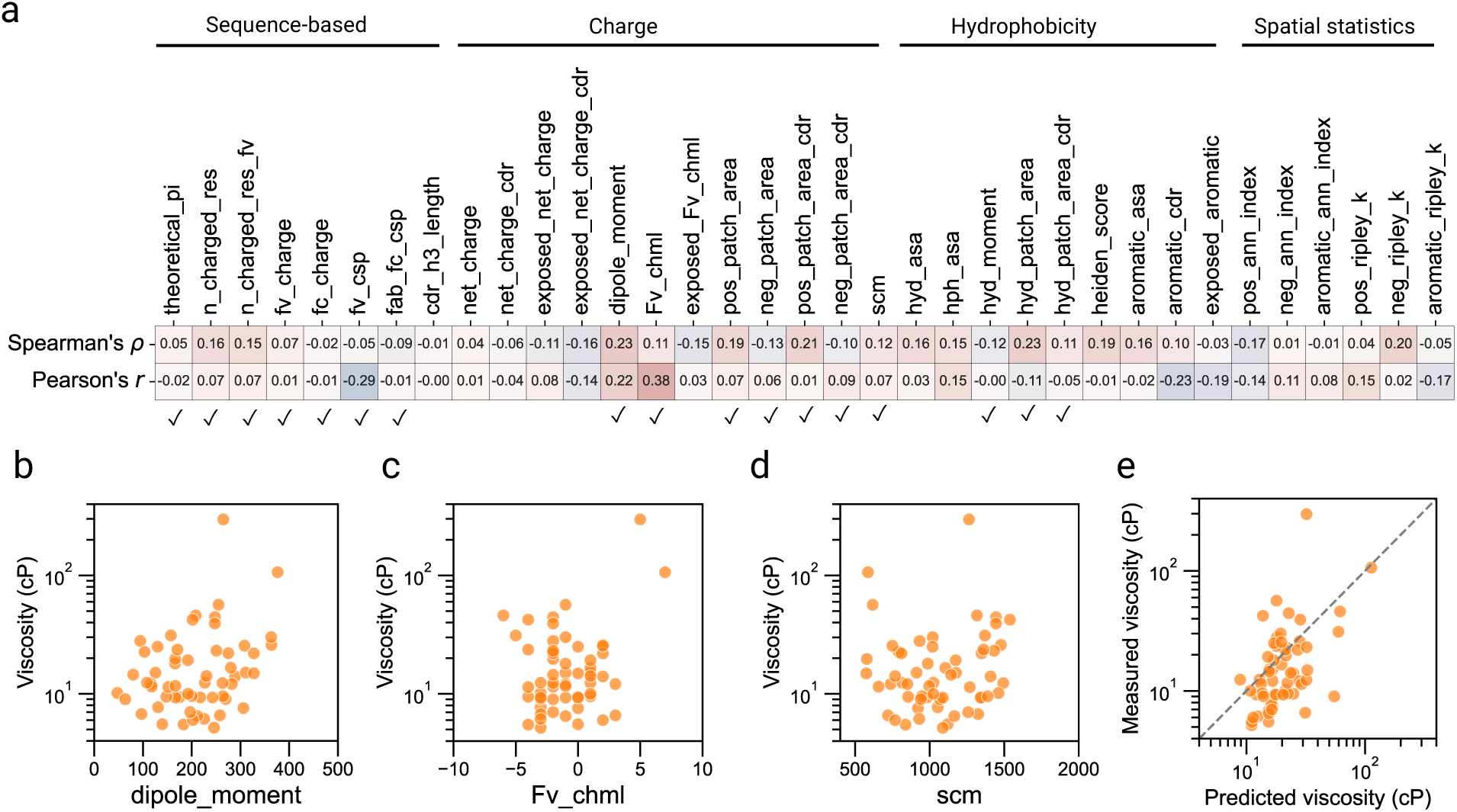
Applying P_ROPERMAB_ to predict high-concentration viscosity of IgG4 mAbs. **a**) Spearman’s ρ and Pearson’s *r* correlation coefficients of each feature with IgG4 mAb viscosity. While there was no single feature that is strongly correlated with viscosity, several charge asymmetry related features such as dipole_moment and Fv_chml demonstrate at least weak correlation with viscosity. **b, c**) Scatter plot of viscosity vs. dipole_moment and Fv_chml respectively, with both having at least some weak correlation with viscosity. **d**) Scatter plot of viscosity vs. the scm feature, showing almost no correlation between the two. **e**) Leave-one-out cross-validation results of a random forest regressor trained to predict viscosity using a set of features manually selected based on in-house expertise (indicated with a ✓ symbol in panel **a**).

We next sought to develop a ML model to predict mAb viscosity. Based on the understanding that hydrophobic and electrostatic interactions are two major contributing factors of high viscosity, we selected a set of hydrophobicity and charge related features as model input. We note that the selection of features was done prior to examining their correlations with viscosity to avoid information leakage [54] and no algorithmic feature selection was done due to the limited training data available. As an example, we trained a random forest model because of its ability in modeling highly non-linear relationships. We followed the same model building and nested cross-validation evaluation procedure as in HIC RT prediction (Methods). As shown in Figure 4e, the random forest model combining multiple features of diverse biophysical nature as input achieved a Spearman’s ρ = 0.48 and a Pearson’s *r* = 0.35. Together with the example on HIC RT prediction, our results demonstrate a broad applicability of the P_ROPERMAB_ framework to *in silico* prediction of mAb developability properties.

### Computational efficiency enables large-scale assessment

Calculating structure-derived features given mAb Fv sequences with P_ROPERMAB_ takes about half a minute on average on an AWS 8-core node with 30GB RAM (Figure 5a). Our calculation suggests that the time and cost associated with calculating the features for a typical secondary screening dataset (∼1000 sequences) is fast and cost-effective (8.5 hours with $0.384/hour). While the computing time scales only linearly with the number of mAb Fv sequences (Supplementary Figure 3), the time and computational cost could be prohibitive if one were to apply P_ROPERMAB_ to large repertoire sequence projects where the scale of sequences can reach millions. We reasoned that if the structure-derived features can be predicted directly from sequence using simple lightweight ML models, then a large fraction of the computing cost in structure prediction and feature calculation can be saved (Figure 5b). As examples, we trained ElasticNet models to predict all structure-derived features from sequences alone. We randomly sampled 12,000 mAbs from the OAS database [55] and divided them into a training set of 10,000 mAbs and a test set with 2,000 mAbs. For each of the mAbs, we calculated the structure-based features using P_ROPERMAB_ and one-hot encoded the sequences after aligning them according to the IMGT numbering scheme [56] (Figure 5b). (Calculating features for a set of 12,000 mAbs would incur one-time cost.) Alignment of sequences to the IMGT numbering scheme [56] enables one-hot encoding with fixed size and biologically meaningful method of padding. Our results show that the structure-derived features can be predicted from sequences alone with high accuracies, with a median Pearson’s r of 0.87 (Figure 5c and Supplementary Figure 4). Encouraged by these results we applied the trained ElasticNet models to predict the structure-based features for the HIC RT and viscosity datasets. While models trained with the predicted features as input experience a slight drop in performance (Figure 5d) compared to models trained with features calculated directly from structures (Figures 3e and 4e), predicting features from sequence is considerably faster than calculating features from structures. In fact, on a MacBook Pro laptop with 16GB RAM and the M1 Pro processor it took less than 2 minutes to predict all 27 structure-based features for >140k paired sequences from the OAS database once the sequences have been one-hot encoded. On the same computing platform, numbering the >140k sequences according to the IMGT scheme and one-hot encoding them took less than 3 hours. Thus, given the accuracy and efficiency of sequence-based prediction, it is feasible to apply P_ROPERMAB_ to large repertoire sequence datasets.

**Figure 5.**
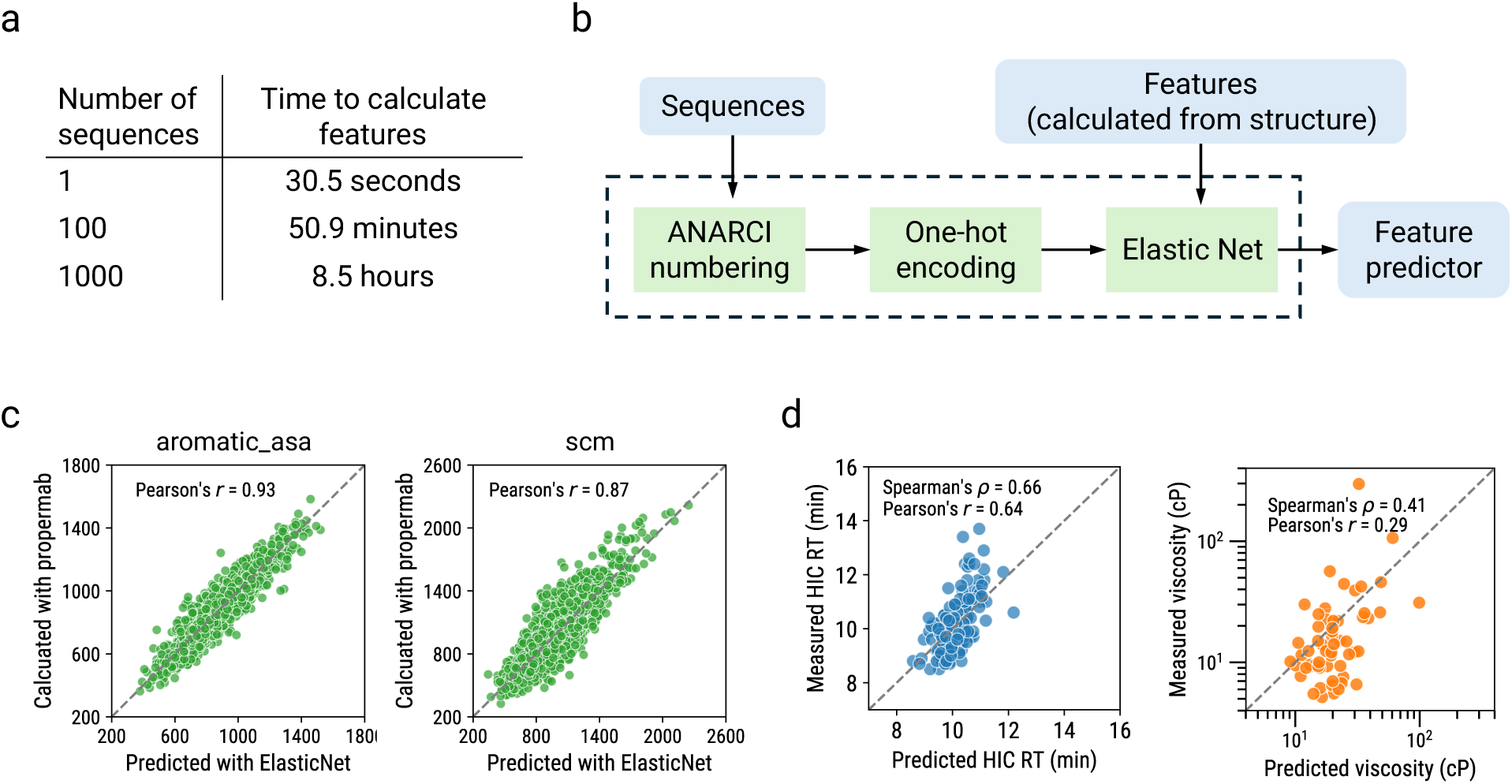
P_ROPERMAB_ enables efficient large-scale feature prediction from sequences. **a**) Time cost of calculating structure-based features using P_ROPERMAB_. We ran and timed P_ROPERMAB_ on 1000 paired sequences randomly sampled from the OAS database [55]. The time for one sequence is the average across 1000 sequences. The time for 100 sequences is the average across a 1000 bootstrap samples. Each bootstrap sample was obtained by randomly sampling 100 individual sequences from the 1000-sequence run. **b**) A schematic showing our strategy to train ElasticNet models to predict structure-based features from sequences. **c**) Comparison of feature values calculated based on structures with values predicted from sequences using trained ElasticNet models for aromatic_asa and scm respectively. The plots are based on a test set of 2000 paired sequences randomly sampled from the OAS database [55]. **d**) Comparison of measured values of HIC RT and viscosity with values predicted using models trained with predicted features as input.

## Discussion

Screening candidates for lead molecules with better developability properties early in the discovery and development of antibody therapeutics has increasingly been recognized as important for success [7, 11, 13, 15]. While multiple biophysical assays have been developed over the years [47], ML models integrating computational molecular features and trained on experimental data could offer a cost-effective approach [13]. We developed P_ROPERMAB_ as an integrative framework with the capability of calculating diverse molecular features and enabling downstream ML model development using these features.

While P_ROPERMAB_ can be applied to a broad spectrum of developability properties, we highlighted its applicability with two use cases. In the case of HIC RT prediction, we showed that several features calculated by P_ROPERMAB_ demonstrated moderate to strong correlation with HIC RT. When an ElasticNet model was trained, it resulted in much higher performance even though about one third of the features were eliminated (i.e. zero model coefficient). The elimination of several features is not unexpected given that some of them are not significantly correlated with HIC RT and that there is some redundancy in the feature space (Supplementary Figure 5). In the case of predicting high-concentration viscosity, we showed that several individual features are highly correlated with viscosity in two previously published datasets of the IgG1 isotype, but no features had strong correlation with viscosity in our IgG4 dataset. Compared to IgG1 mAbs, there is much less viscosity data of IgG4 mAbs. Our finding based on a relatively large internal IgG4 dataset suggests that the mechanisms underlying high viscosity of IgG4 mAbs is likely more complex than IgG1 mAbs and that engineering a single, strongly predictive feature requires better understanding of the underlying biophysical mechanisms. Nevertheless, when multiple features were integrated into a ML model, prediction of IgG4 viscosity is improved. We note that in the current work the predictive power of individual features is evaluated based on their linear correlations, which can fail to quantify associations that are nonlinear [57]. A possible future development in feature selection is to employ an information-theoretic formalism in contrast to the traditional correlation-based approaches. For example, by quantifying the mutual information, a self-equitable measure of association, between features and the biophysical properties, one could select features that maximize information gain and thereby improve prediction [57].

Recently, there has been interest in applying *in silico* developability assessment tools to large repertoire sequence datasets that may contain ∼10^10^ sequences. While structure-derived features are critical for antibody developability prediction [58], calculating the structure-derived features for each molecule at this scale is computationally prohibitive. Here, we showed that structure-derived features as calculated using P_ROPERMAB_ can be predicted with high accuracy from sequence alone with relatively simple models. Previously, deep learning surrogate models were trained to predict mAb spatial properties such as the SCM score with target features in the training sets calculated as ensemble averages from 10-ns long molecular dynamics (MD) trajectories [59, 60]. While computationally more expensive structure modeling such as MD simulations may generate important mechanistic insights, recent results on its benefit to ML-based developability prediction are not yet consistent [20, 22]. In this work, we demonstrated a computationally much more efficient and practically effective alternative to predicting structure-based features from sequences only. We trained simpler ElasticNet models to predict features calculated by P_ROPERMAB_ and avoided the use of expensive MD simulations. Importantly, we showed that ML models using ElasticNet predicted features as input resulted in only a slight drop in performance. The efficiency and effectiveness of our method suggest that it can be a viable approach for *in silico* developability prediction of repertoire-scale datasets.

Additionally, it was recently shown there is poor agreement between several structure-derived features calculated using different software packages or different versions and protocols of the same software, highlighting the need for transparency in algorithmic implementations and sharing of the structures used to aid reproducibility and comparison [14]. While some of the features (for example, the patch related features) calculated by P_ROPERMAB_ can be calculated using other software packages, the lack of access to the implementation details of these features in other software packages hinders reproducibility and comparison. This is further amplified by the broad spectrum of developability properties usually considered in antibody development and the potentially infinite number of molecular features that can be calculated from sequences and structures. We believe that the open-source nature of P_ROPERMAB_ will aid the comparison and development of new molecular features. One future direction for developing the package would be to represent antibody structures as molecular graphs to leverage the latest geometric deep learning (GDL) architectures for molecular property prediction [61–63]. While P_ROPERMAB_’s current Sequence Featurizer component calculates features based on full sequences, the Structure Featurizer module only calculates features derived from the Fv domain, as current structure prediction methods for immune proteins such as ABodyBuilder2 [36] only predicts the 3D structure of the Fv domain. Whole-mAb structures and features derived from them are expected to further improve *in silico* prediction of developability properties.

While having the capability to calculate informative molecular features at large scale with low cost is critical, a major bottleneck in developing accurate ML models for developability prediction has been the lack of large and high-quality training data. Depending on the developability property being studied, the total number of datapoints in a training set can be as low as a few dozen (for example, viscosity) [25, 26]. The issue of lack of training data is further complicated by discrepancies in experimental data such as unmatched experimental conditions, different measuring instrumentations, and batch effects. Training models using aggregated datasets or applying models across datasets can easily violate the implicit assumption underlying many ML algorithms that the training data are independently and identically distributed. Such violations may partially explain the poor transferability of some recently developed developability prediction models [14, 25]. Thus, targeted generation of large-scale experimental measurements under consistent and well controlled conditions is desirable and expected to alleviate these issues.

## Methods

### Calculating sequence-based features

The theoretical_pi feature is calculated based on the Henderson-Hasselbach equation by accounting for six amino acids D, E, H, K, R, and Y with ionizable side chains, and mAb terminal groups [64]. According to the Henderson-Hasselbach equation, the net charge *q* of a mAb at a given pH is

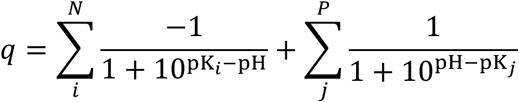

where *i* runs through all negatively charged amino acids (including tyrosine), *j* runs through all positively charged amino acids, *N* and *P* are the number of negatively and positively charged amino acids respectively. To determine the theoretical pI, we consider the full mAb sequence, i.e. a pair of heavy chains and a pair of light chains, and the theoretical pI is determined as the pH that minimizes the square of the net charge *q*. For amino acid pK values, we used the pK set available at EMBOSS [65].

We calculate n_charged_res by counting the total number of charged residues in full mAb sequence. At the default pH = 7.4, we consider residues D, E, K, R as charged. Similarly, n_charged_res_fv is computed by counting the total number of charged residues in the Fv domain. fv_charge is computed as the total charge of the Fv domain. At the default pH = 7.4, we assign residues D and E a −1 charge, and K and R a +1 charge. fv_csp stands for the charge separation between the VH domain and VL domain and is defined as the total charge of the VH domain times the total charge of the VL domain. Similarly, fab_fc_csp stands for the charge separation between the Fab domain and the Fc domain and is defined as the total charge of the Fab domain times the total charge of the Fc domain. We define the Fab-Fc boundaries based on UniProt sequence annotations [66]. cdr_h3_length stands for the length of the CDR-H3 loop and is calculated by counting the number of residues in the CDR-H3 loop as defined according to the IMGT numbering scheme [56].

### Calculating structure-based features

All structure-based features are calculated based on 3D structures of the Fv domain that is numbered according to the IMGT numbering scheme [56]. 3D structures of Fv domains are predicted using the antibody specific ABodyBuilder2 tool of the ImmuneBuilder suite (version 0.0.8) [36]. In general, we assigned atomic partial charges according to the CHARMM36 force field [67] using the molecular system and topology utilities from the OpenMM Python API (version 8.0) [68]. Solvent accessibility of atoms and residues are computed using the FreeSASA package (version 2.1.0) [69]. An atom is considered solvent exposed if its solvent accessibility is greater than 0, and a residue is considered solvent exposed if its relative solvent accessibility (RSA) is greater than or equal to 0.05. To calculate patch-related features, we first used NanoShaper (version 0.7.8) [70] to calculate a triangulated mesh representation of the molecular surface of the Fv structure with grid_scale = 0.5. We then used the APBS tool (version 3.0.0) [71] to calculate the Poisson-Boltzmann electrostatic potentials and the multivalue utility available as part of APBS tool to assign electrostatic potential at each vertex of the meshed surface. To assign hydrophobic potentials to mesh vertices, we implemented the algorithm by Heiden et al. [72, 73]. In the following, we describe the algorithms we implemented to calculate individual structure-based features.

#### net_charge and exposed_net_charge

The net_charge feature of the Fv domain is calculated by summing up the partial charges of all atoms in the domain. Similarly, the exposed_net_charge feature is calculated by summing up the partial charges of all solvent-exposed atoms.

#### net_charge_cdr and exposed_net_charge_cdr

The net_charge_cdr feature is the total charge in CDR regions. It is calculated by summing up the partial charges of all atoms located in CDR regions. The exposed_net_charge_cdr feature is the total charge of CDR atoms that are solvent-exposed and is calculated similarly.

#### scm

The scm feature stands for the spatial charge map score. In P_ROPERMAB_ we calculate the scm according to the algorithm described in [29].

#### Fv_chml and exposed_Fv_chml

The Fv_chml is a feature that describes the charge asymmetry between the VH domain and the VL domain. It is calculated by subtracting the net charge of the VL domain from the net charge of the VH domain. The exposed Fv_chml feature describes the surface charge asymmetry between the VH domain and VL domain. It is calculated by subtracting the exposed net charge of the VL domain from that of the VH domain.

#### dipole_moment

Calculating the dipole moment of a protein is a complex topic. However, because the dipole moment of a charge distribution is always defined as

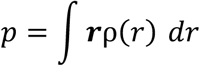

where ρ(r) is the charge density at location r. If the charge distribution is a collection of point charges q_i_ at location r_i_ then the integral simplifies to a summation

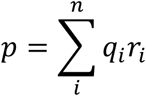

In P_ROPERMAB_ we treat the charge distribution of the Fv domain as a collection of point charges at atom locations and use the summation above to compute the dipole moment of the Fv domain. The locations *r*_i_ are centered at the center of geometry of the domain, that is r_i_ = r_i0_ − r_0_, where r_i0_ is the original coordinates in the PDB file, and

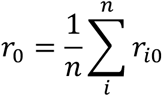

is the center of geometry of the domain. We note that the dipole moment is multiplied by 4.803 to convert from Angstrom-electron-charge units to Debyes as described in [74].

#### aromatic_cdr and exposed_aromatic

This aromatic_cdr feature is calculated by counting the total number of aromatic residues (i.e. F, W, Y) across all CDR regions, which are defined according to the IMGT numbering scheme as previously described. The exposed_aromatic feature is calculated by counting the total number of aromatic residues in the Fv domain that are solvent exposed. A residue is solvent exposed if its RSA is 0.05 or bigger as previously described.

#### hyd_moment

The hyd_moment feature stands for hydrophobic moment, which is the analogue of the electric dipole moment for hydrophobicity. Just as the electric dipole moment measures the asymmetry of the charge distribution, the hydrophobic moment measures the amphiphilicity (asymmetry of hydrophobicity) of the structure. In P_ROPERMAB_ the hydrophobic moment is computed as

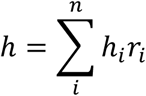

where h_i_ is the hydrophobicity of residue *i* and r_i_ is the center of geometry of residue *i*. P_ROPERMAB_ offers multiple options for amino acid hydrophobicity scales, the Kyle-Doolittle (KD) scale [75] is used by default.

#### heiden_score

The heiden_score feature is calculated by summing up the hydrophobic potentials of all surface vertices whose hydrophobic potential is positive as described in [73]. The hydrophobic potential of each vertex is weighted by its surface area, which is calculated by splitting the area of each triangle evenly to the three vertices. The hydrophobicity potential of individual vertex is a weighted average of the lipophilicities of its neighboring atoms as described in [73] and [72]. For atomic lipophilicity scale, we use the parameters described in [76].

#### hyd_asa, hph_asa, and aromatic_asa

The hyd_asa feature stands for the hydrophobic solvent accessible surface area. It is calculated by summing up the apolar component of the solvent accessible surface area of all residues of the Fv domain. Similarly, the hydrophilic solvent accessible surface area (hph_asa) is calculated by summing up the polar component of the solvent accessible surface area of all residues. Calculation of solvent accessible surface areas and their decomposition into polar and apolar components are done using FreeSASA, as previously described. The total solvent accessible aromatic surface area (aromatic_asa) is calculated by summing up the solvent accessible surface areas of aromatic residues.

#### pos_ann_index, neg_ann_index, and aromatic_ann_index

For a given set of residues, the Average Nearest Neighbor (ANN) metric measures the average distance between the residues and their nearest neighbors. If the average distance is less than the average for a hypothetical random distribution, then the distribution of the residues being analyzed is considered clustered. Otherwise, the residues are considered dispersed. In P_ROPERMAB_ we calculate the ratio of the observed average distance to the expected average distance and name this feature ann_index, i.e.

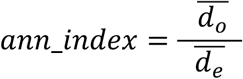

where ^—^d_o_ is the average of observed distance between each residue and its nearest neighbor:

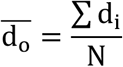

and ^—^d_e_ is the expected average distance for the set of residues under a null distribution and *N* is the size of the set. We simulate the null distribution by permuting the residues over locations of all solvent-exposed residues. Currently, P_ROPERMAB_ implements this feature for positively charged (pos_ann_index), negative charged (neg_ann_index), and aromatic residues (aromatic_ann_index), respectively.

#### pos_ripley_k, neg_ripley_k, and aromatic_ripley_k

The Ripley’s K function is a spatial statistic that summarizes spatial dependence (residue clustering or dispersion) over a range of distances. For a given set of residues, the Ripley’s K calculates the average number of neighboring residues associated with each residue at a given distance cutoff. If the average number of neighbors at the distance cutoff is higher than the average number of neighbors expected under a null distribution, then the residues are considered clustered. In P_ROPERMAB_, the Ripley’s K statistic (named ripley_k) is defined as

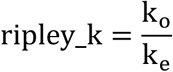

where k_o_ is the observed proportion of neighboring residues evaluated at the given distance cutoff d_cutoff_, i.e.

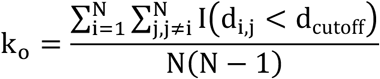

where *N* is the total number of residues, d_i,j_ is the Euclidean distance between residues *i* and *j* and *I* is the indicator function that evaluates to 1 when *d*_*i,j*_ is less than *d*_cutoff_ and 0 otherwise. k_e_is the expected proportion of neighboring residues for the evaluated distance under a null distribution. We simulate the null distribution by permuting the residues over the locations of all solvent-exposed residues. Currently, P_ROPERMAB_ implements this feature for positively charged (pos_ripley_k), negative charged (neg_ripley_k), and aromatic residues (aromatic_ripley_k), respectively.

#### hyd_patch_area, hyd_patch_area_cdr, pos_patch_area, pos_patch_area_cdr, neg_patch_area, neg_patch_area_cdr

To detect surface patches of a given biophysical property (positively charged, negatively charged, or hydrophobic), we first assign a property value to each triangle on the surface mesh by taking the average value of its three vertices. We then remove all triangles whose property value does not meet a given property threshold. For the remaining triangles, we employ the DBSCAN algorithm [77] as implemented in scikit-learn to cluster them based on the Cartesian coordinates of their geometric centers. Clusters of triangles whose total areas exceed a given area threshold are considered as surface patches. We set the area thresholds on positively charged and negatively charged patches at 20 Å^2^, and the threshold on hydrophobic patches at 40 Å^2^, determined through empirical evaluation and visual inspection. A patch is considered near CDR if there is at least one vertex of the patch that is within 5 Å of any CDR vertices. NanoShaper assigns vertices to residues, and we define CDR vertices as the set of vertices that get assigned to the residues of CDR regions.

#### Sequence embeddings

Protein language models (PLMs) have seen wide applications in protein property prediction and design [78–80]. PLMs transform textual protein sequences into meaningful high-dimensional numerical representations called sequence embeddings that can be fed as input to downstream machine learning tasks. While not discussed in detail in the current work, we implemented the SeqEmbedder class in P_ROPERMAB_ to facilitate the embedding of sequences using the ESM-1b [34] and AntiBERTy [37] models. Embeddings from these models have recently been shown to be predictive of antibody developability properties such as solubility [16] and thermostability [21].

#### 3D voxel grids

The 3D structures of proteins can be treated as if they were multi-channel 3D images known as 3D voxel grids [81]. Such representation makes protein structures amenable to 3D convolutional neural networks (3D-CNNs) and has enabled the application of 3D-CNNs in various protein related tasks such as protein-ligand binding affinity prediction [82], prediction of changes in thermodynamic stability [83], protein design [84], and more recently, prediction of antibody viscosity [26]. However, creating 3D voxel grid representations of protein structures can be a challenging task. In P_ROPERMAB_, we implemented a simple interface that enables the transformation of protein 3D structure coordinates into 3D voxel grid representations with a single function call, while hiding the details of voxel creation and featurization. Such functionality and simple interface are expected to facilitate the use of 3D-CNNs for antibody developability prediction.

#### Training and evaluating ML models for predicting HIC RT and viscosity

We trained and evaluated the ML model as implemented in the scikit-learn package (version 1.2.2). Our model building procedure consists of wrapping feature standardization via a StandardScaler and parameter fitting of the ElasticNet model in a pipeline and using GridSearchCV to fit parameters and tune hyperparameters of the model in the pipeline. As an example, we searched a combination of the l1_ratio and the alpha parameters, with the l1_ratio ranging from 0.1 to 0.9 with a step size of 0.1, and four alpha values of 0.01, 0.05, 0.1, and 0.5. We then nested our GridSearchCV procedure within a leave-one-out cross-validation loop to estimate the generalization performance of our model building procedure. Such a nested cross-validation is necessary to avoid overestimation of model generalization ability [85]. The RandomForestRegressor model for viscosity prediction is trained and evaluated by following the same procedure as HIC RT prediction.

## Data availability

The molecular features computed using P_ROPERMAB_ for the HIC RT dataset and the Ab21 and PDGF38 viscosity datasets are available as Supplementary Datasets.

## Code availability

The source code of P_ROPERMAB_ is available at https://github.com/regeneron-mpds/PROPERMAB for non-commercial uses.

## Acknowledgement

The authors would like to thank Robert Babb and Calvin Chen from Regeneron Protein Expression Sciences for their discussions and suggestions during the project. The authors also thank Eric Prager from Regeneron Research Program Management for his administrative support.

