## Supplementary Information for "PROPERMAB: an integrative framework for *in silico* prediction of antibody developability using machine learning"

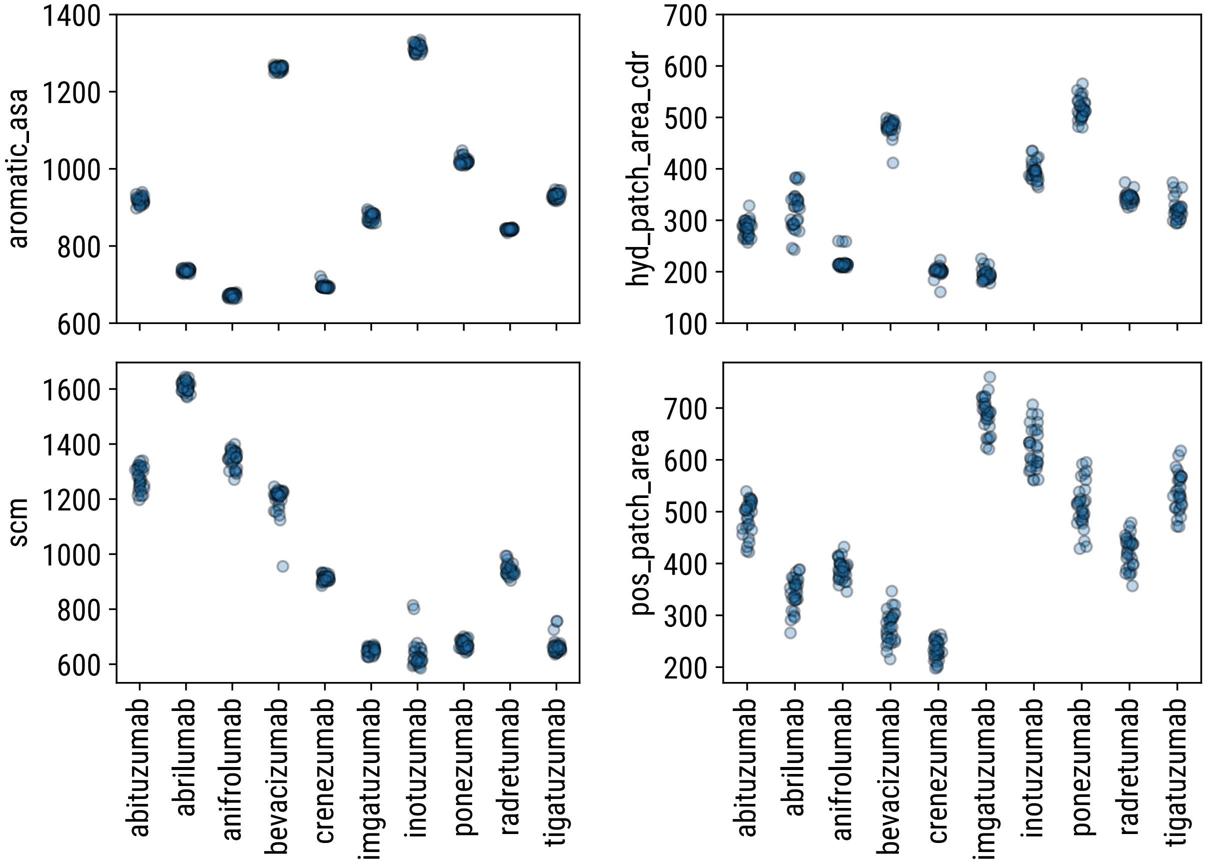


**Supplementary Figure 1.** Examples of feature variability. Values of structure-based features can vary across runs to various degrees due to the randomness in the structure relaxation of ImmuneBuilder [1]. Here we ran our feature pipeline 30 times for each molecule to explore the variability across runs. Empirically, we found 5 runs to be a good compromise between mitigating variability and computational cost.

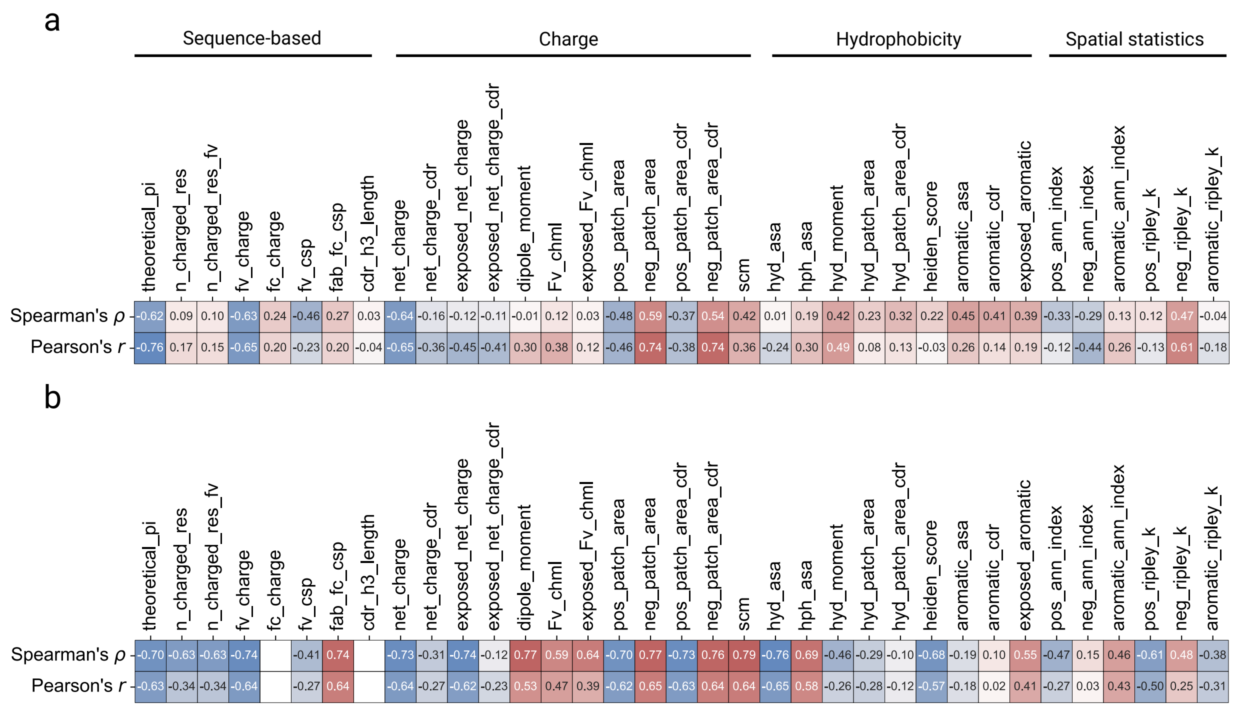


**Supplementary Figure 2.** Correlations of propermab features with viscosities from two previously published datasets, namely Ab21 a) and PDGF38 b). For the PDGF38 dataset, the values of fc_charge and cdr_h3_length are constant because all molecules are derived from the same parent molecule through single-point mutations. Thus, the correlations of fc_charge and cdr_h3_length with viscosity cannot be calculated.

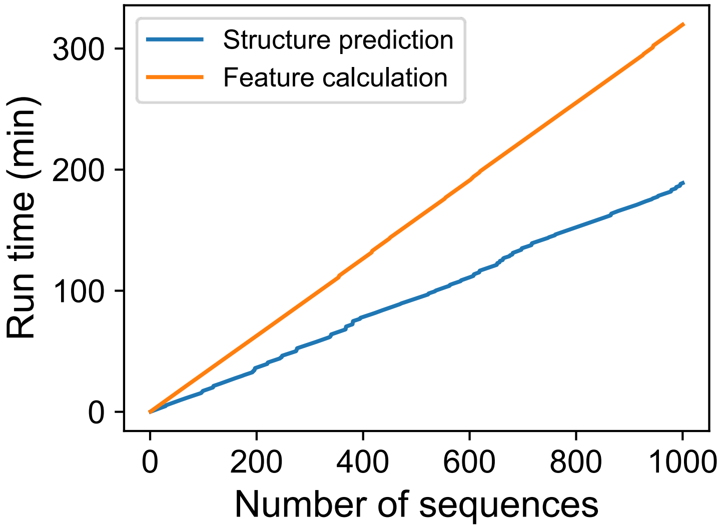


**Supplementary Figure 3.** Assessment of run time of the propermab feature pipeline. The propermab feature pipeline consists of 3D structure prediction using ImmuneBuilder [1] and calculation of sequence-based features and 3D structure-based features. As shown here, both structure prediction and feature calculation scales linearly with the number of sequences.

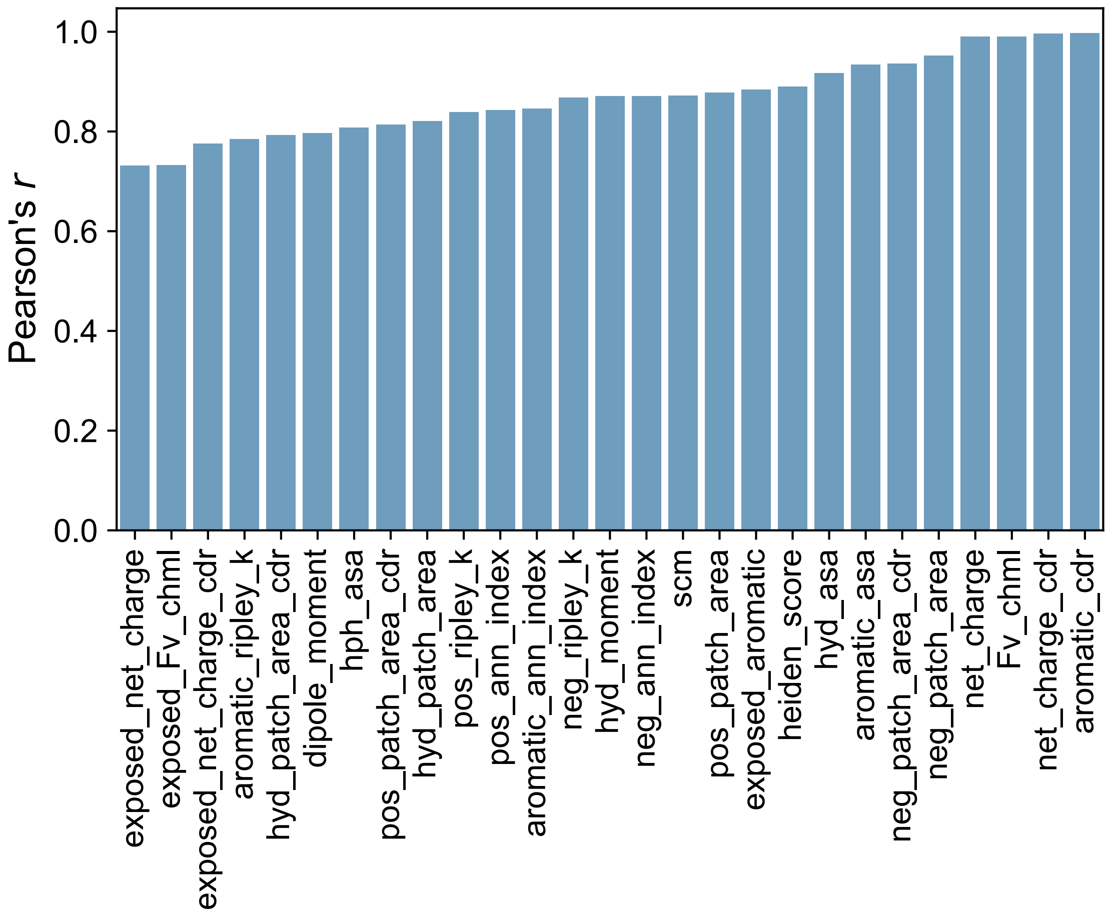


**Supplementary Figure 4.** Performance of ElasticNet models on predicting structure-based features from onehot-encoded sequences. The plots are based on a test set of 2000 paired sequences randomly sampled from the OAS database [2].

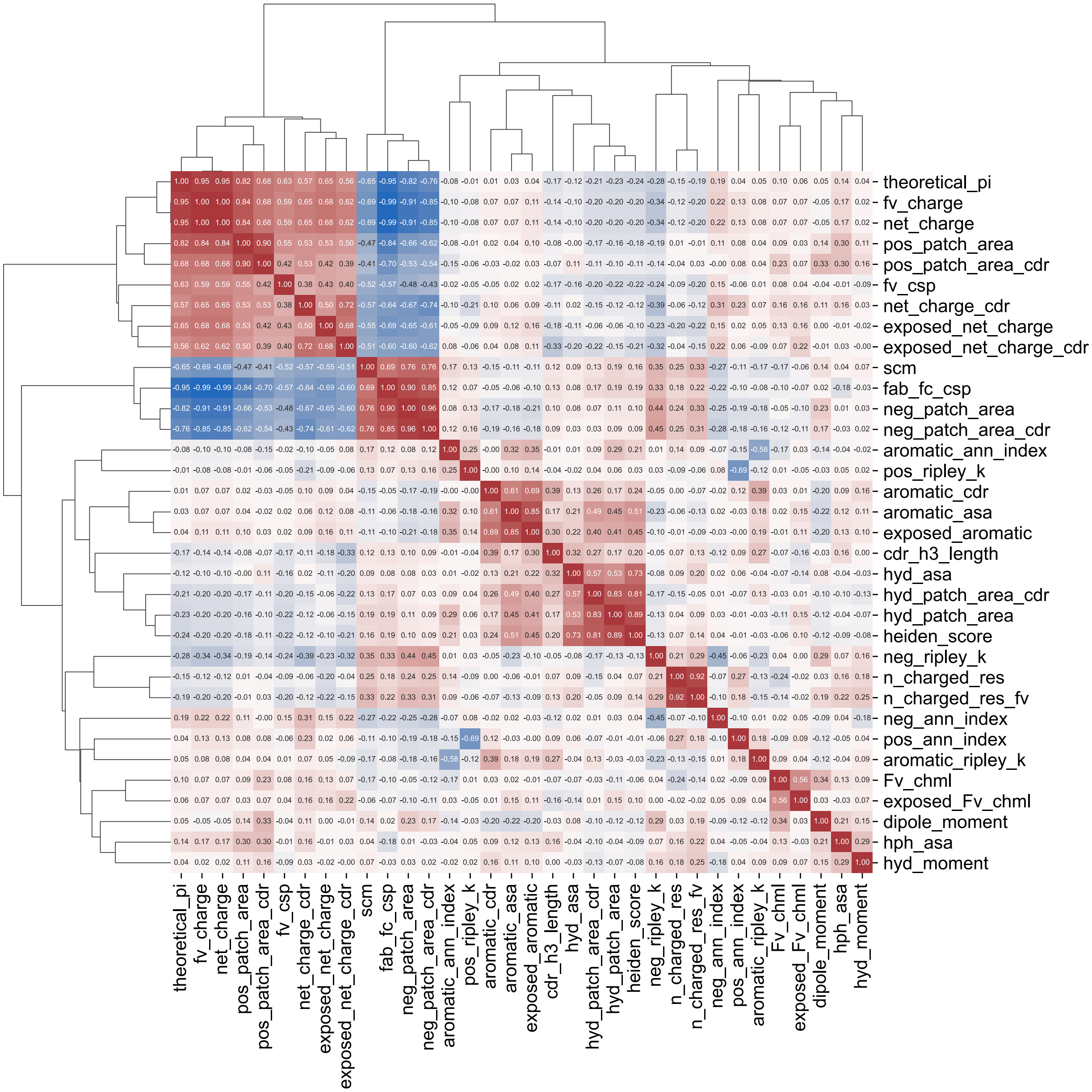


**Supplementary Figure 5.** Pearson correlations between pairs of features. The correlations were calculated based on features for 135 molecules from [3].

**Supplementary Table 1.** Summary of sequence-based features

| Feature name | Brief description |
|---|---|
| theoretical_pi | Theoretical isoelectric point. |
| n_charged_res | Total number of charged residues. |
| n_charged_res_fv | Total number of charged residues in the variable domain. |
| fv_charge | Charge of the variable domain. |
| fc_charge | Charge of the constant domain. |
| fv_csp | Charge separation of the variable domain, i.e. charge of the VH domain times the charge of the VL domain. |
| fab_fc_csp | Charge separation of the Fab domain and constant domain, i.e. charge of the Fab domain times the charge of the constant domain. |
| cdr_h3_length | Number of residues in the CDR-H3 loop, as defined according to the IMGT numbering scheme. |

Details of the algorithms for calculating these features are described in the Methods section.

**Supplementary Table 2.** Summary of structure-based features.

| Feature group | Feature name | Brief description |
|---|---|---|
| Charge related | net_charge | Net charge of the Fv domain. |
| | net_charge_cdr | Net charge of CDR regions. |
| | exposed_net_charge | Net charge of solvent-exposed atoms. |
| | exposed_net_charge_cdr | Net charge of solvent-exposed atoms in CDR regions. |
| | dipole_moment | The electric dipole moment of the Fv domain. |
| | Fv_chml | The charge asymmetry between the VH domain and the VL domain. |
| | exposed_Fv_chml | The charge asymmetry between the VH domain and the VL domain calculated based on atoms that are solvent-exposed. |
| | pos_patch_area | Total area of positively charged surface patches. |
| | pos_patch_area_cdr | Total area of positively charged surface patches that are located near CDR regions. |
| | neg_patch_area | Total area of negatively charged surface patches. |
| | neg_patch_area_cdr | Total area of negatively charged surface patches that are located near CDR regions. |
| | scm | The spatial charge map. |
| Hydrophobicity related | hyd_asa | Total hydrophobic solvent accessible surface area of the Fv domain. |
| | hph_asa | Total hydrophilic solvent accessible surface area of the Fv domain. |
| | hyd_moment | The dipole moment analogue for hydrophobicity. |
| | hyd_patch_area | Total area of hydrophobic patches. |
| | hyd_patch_area_cdr | Total area of hydrophobic patches that are located near CDR regions. |
| | heiden_score | The Heiden hydrophobic score for the Fv domain. |
| | aromatic_asa | Total solvent accessible surface area of aromatic residues. |
| | aromatic_cdr | Total number of aromatic residues in CDR regions. |
| | exposed_aromatic | Total number of solvent-exposed aromatic residues. |

| | | |
|---|---|---|
| Spatial statistics | pos_ann_index | The average nearest neighbor index of solvent-exposed, positively charged residues. |
| | neg_ann_index | The average nearest neighbor index of solvent-exposed, negatively charged residues. |
| | aromatic_ann_index | The average nearest neighbor index of solvent-exposed, aromatic residues. |
| | pos_ripley_k | A variant of the Ripley's K statistic of solvent-exposed, positively charged residues. |
| | neg_ripley_k | A variant of the Ripley's K statistic of solvent-exposed, negatively charged residues. |
| | aromatic_ripley_k | A variant of the Ripley's K statistic of solvent-exposed, aromatic residues. |

Details of the algorithms for calculating these features are described in the Methods section.

**Supplementary Table 3.** Mean value of pos_patch_area across multiple runs.

| Molecule | 1_run | 5_run | 10_run | 20_run | 30_run |
|---|---|---|---|---|---|
| abituzumab | 474.2 | 504.8 | 489.1 | 493.4 | 489.8 |
| abrilumab | 266.1 | 325.6 | 342.6 | 337.5 | 342.9 |
| anifrolumab | 364.6 | 385.7 | 387.8 | 387.9 | 387.7 |
| bevacizumab | 215.5 | 274.2 | 276.9 | 275.5 | 277.5 |
| crenezumab | 251.8 | 245.5 | 237.5 | 235.8 | 233.0 |
| imgatuzumab | 690.9 | 705.9 | 710.3 | 694.5 | 687.1 |
| inotuzumab | 594.9 | 600.2 | 608.8 | 615.6 | 620.9 |
| ponezumab | 497.7 | 532.8 | 525.4 | 506.2 | 511.5 |
| radretumab | 399.1 | 427.8 | 428.1 | 427.1 | 422.8 |
| tigatuzumab | 507.3 | 542.3 | 540.9 | 526.2 | 537.2 |
